# Leukocytes have a heparan sulfate glycocalyx that regulates recruitment during inflammation

**DOI:** 10.1101/2024.05.21.595098

**Authors:** Megan J. Priestley, Anna K. Hains, Iashia Z. Mulholland, Sam Spijkers-Shaw, Olga V. Zubkova, Amy E. Saunders, Douglas P. Dyer

## Abstract

The glycocalyx is a proteoglycan-rich layer present on the surface of all mammalian cells that is particularly prevalent on endothelial cells lining the vasculature. It has been hypothesized that the glycocalyx mediates leukocyte migration by masking adhesion molecules and reducing leukocyte adhesion to the endothelium. Leukocyte recruitment is a key driver of inflammatory diseases, including the chronic skin disease, psoriasis. Here, we show that leukocytes express heparan sulfate, an important glycocalyx component, on their cell surface which is lost in response to psoriasis-like skin inflammation, whilst endothelial heparan sulfate expression is not affected. Treatment with a heparan sulfate mimetic during psoriasis-like skin inflammation protected heparan sulfate from cleavage by heparanase and resulted in reduced leukocyte accumulation in skin, yet unexpectedly, led to increased clinical signs of inflammation due to reduced Treg numbers. These findings reshape our understanding of immune cell recruitment by revealing the presence and function of a heparan sulfate glycocalyx on immune cells and highlight the complex effects of heparanase inhibitors on the immune response in this context.

**One Sentence Summary:** Leukocytes express a glycocalyx on their surface which is shed in response to psoriasis-like skin inflammation, facilitating their migration into the skin.

## INTRODUCTION

During inflammatory disease, excessive recruitment of leukocytes results in aberrant inflammation^1^. The chronic inflammatory skin disease, psoriasis, is associated with the accumulation of large numbers of leukocytes in the dermis, resulting in raised, itchy, and scaly skin lesions^2^. Despite recruitment of immune cells into tissues being a clear driver of inflammation, there are currently no therapeutic strategies targeting leukocyte recruitment as a treatment for psoriasis^3^. In fact, although studied for many years, there are still large gaps in knowledge about the process of leukocyte migration in general^4^. Much research on leukocyte transmigration out of blood vessels has focussed on endothelial cell adhesion molecules such as selectins and immunoglobulin-like adhesion molecules^5^, yet much less attention has been paid to the molecules which initiate and regulate the initial interactions between leukocytes and the endothelium; proteoglycans^6^.

All mammalian cells likely express proteoglycans on their surface to varying extents, as a part of the cell surface glycocalyx^7^. This layer is formed of core proteins such as syndecans and glypicans, and glycosaminoglycans such as heparan sulfate^8^. Despite glycocalyces being present on the surface of all cells, research until now has focussed heavily on the glycocalyx of the vascular endothelium. Here, the glycocalyx protrudes up to 2 microns from the cell surface^9^, making it one of the first layers to come into contact with leukocytes within blood vessels, and potentially regulating early interactions between leukocytes and endothelia^10^. The glycocalyx has also been shown to support chemokine function^11, 12^, altering their availability to leukocytes and thereby affecting migration.

However, the glycocalyx is also thought to have anti-adhesive properties due to its shielding of endothelial cell adhesion molecules from binding by leukocytes^9^. During inflammation, the glycocalyx can be shed by factors including shear stress^13^, reactive oxygen species^14^, and enzymatic degradation by enzymes such as heparanase, which cleaves heparan sulfate from the glycocalyx^15^. This shedding of the endothelial glycocalyx has pro-migratory effects, thought to be due to increased exposure of adhesion molecules which are ‘masked’ by the glycocalyx at rest^10^. Glycocalyx shedding is associated with several inflammatory conditions including sepsis^16^, COVID-19^17^, diabetes^18^ and recently, psoriasis^19^.

Compared to the growing number of studies on the role of the endothelial glycocalyx in inflammatory diseases, very little is known about its counterpart; the leukocyte glycocalyx. Previous studies have shown the presence of proteoglycans, including heparan sulfate proteoglycans, on the surface of leukocytes^20–22^, yet little is known about the structure of this layer or its role in the regulation of leukocyte migration.

Here, we show that a model of psoriasis-like skin inflammation is not associated with endothelial HS shedding, however changes were observed in the leukocyte glycocalyx, which was shown to be present on the surface of skin leukocytes *in vivo*. Furthermore, we demonstrate that the use of the heparan sulfate mimetic Tet-29 reduces immune cell recruitment into skin, potentially via its inhibitory action on the heparanase enzyme. Counterintuitively, this reduced immune cell recruitment exacerbated skin inflammation, which may be due to reduced regulatory T cell recruitment. Our findings highlight a novel mechanism for immune cell recruitment *in vivo*, whereby myeloid cells release heparanase and degrade HS on the surface of leukocytes, facilitating their entry into the tissue.

## RESULTS

### Aldara cream stimulates elevated levels of circulating heparan sulfate but is not associated with endothelial heparan sulfate shedding

During inflammation, leukocytes are recruited into tissues by passing through the vascular endothelium^23^. Therefore, it was hypothesized that the endothelial glycocalyx played a role in psoriasis-like skin inflammation by regulating immune cell recruitment. To assess this, numbers of leukocytes in naïve skin and in psoriasis-like skin inflammation were quantified to determine if immune cells were recruited. Aldara cream was applied topically to the ear pinnae of mice for 6 consecutive days (**Fig 1A**), which led to visible erythema (redness) and scaling of the skin (**Fig 1B**), epidermal thickening **(Fig 1C, D)**, and increased clinical signs of inflammation (**Fig S1A, B, C**) as shown previously^24, 25^. Crucially, Aldara cream induced the accumulation of large numbers of CD45+ cells into skin, peaking at day 6 (**Fig 1E**), with significantly higher numbers of dendritic cells, MHCII+ macrophages, monocytes, neutrophils, and TCRβ+ T cells in Aldara treated skin compared to naive skin (**Fig 1F**), gating shown in **Fig S2**.

**Figure 1.**
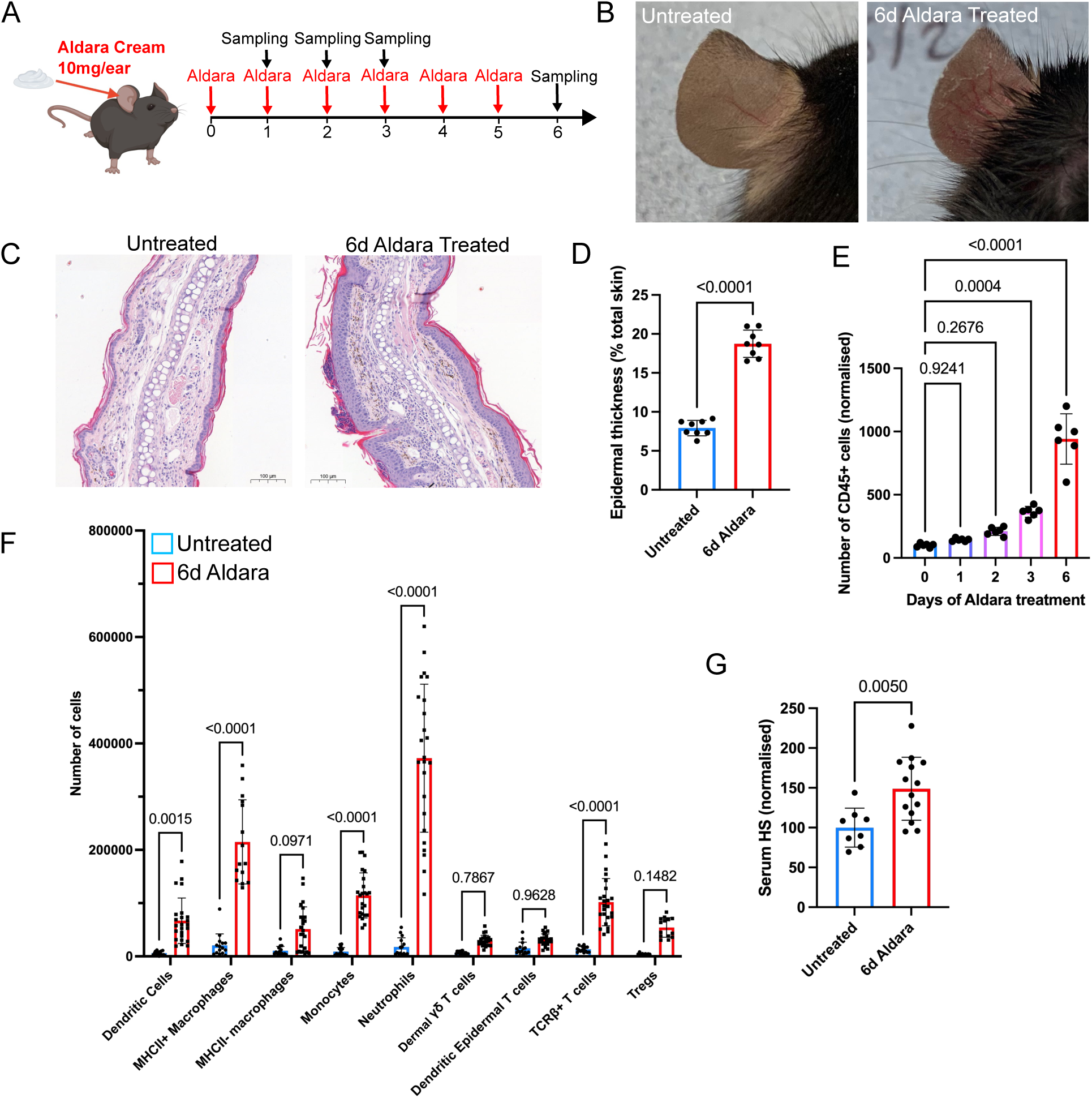
Topical application of Aldara cream stimulates immune cell accumulation in skin and elevates serum heparan sulfate. Mice were treated with topical application of 10 mg Aldara cream to each ear pinnae daily for 0-6 days (**A**) to induce skin inflammation (**B**). Skin was sectioned and stained using haematoxylin and eosin (H&E) (**C**) and epidermal thickness measured from these images and calculated as a % of total ear thickness (**D**). The number of CD45+ cells normalised to the mean of untreated controls (**E**) and numbers of dendritic cells, macrophages, monocytes, neutrophils and T cells (**F**) were quantified on day 6 of Aldara treatment in both ears by flow cytometry. Circulating HS in the blood was measured by ELISA and normalised to the untreated control (**G**). Histology scale bars represent 100 μm. Data are pooled from 3 independent experiments, except for **F** which is pooled from 4 independent experiments. Each point represents individual mouse data. **B and C** show representative images from one experiment, with **C** at 10x magnification. Data in **D and G** were analysed by unpaired t test, **E** by one-way ANOVA with Tukey’s multiple comparisons test and **F** by 2-way ANOVA with Tukey’s multiple comparisons test. Error bars represent mean ± SD.

Psoriasis is associated with enhanced angiogenesis^26^ which may allow increased migration of leukocytes out of blood vessels and into skin^27^. Similarly, Aldara cream treatment resulted in greater skin vascularisation when compared to naïve skin (**Fig S1D, E, F**). To determine whether these vascular changes are accompanied by changes in the endothelial glycocalyx lining the blood vessel, the level of serum HS was measured by ELISA. Aldara cream treated mice had significantly higher concentrations of HS in their serum compared to naïve mice (**Fig 1G**), an indicator of glycocalyx shedding^7,10^.

To determine whether the source of elevated circulating HS was the dermal endothelial glycocalyx, skin sections underwent immunofluorescence staining for HS alongside CD31 as a marker for vascular endothelial cells (**Fig 2A, B**), revealing no significant changes in HS mean fluorescence intensity (MFI) across blood vessels following administration of Aldara cream (**Fig 2C**). HS levels were also similar on naïve and inflamed skin endothelial cells (CD45-CD31+) when examined by flow cytometry (**Fig 2D, E**), although the possibility of ultrastructural changes cannot be excluded. These data indicate that although heparan sulfate is elevated in the circulation of mice with psoriasis-like skin inflammation, the source does not appear to be the endothelial glycocalyx.

**Figure 2.**
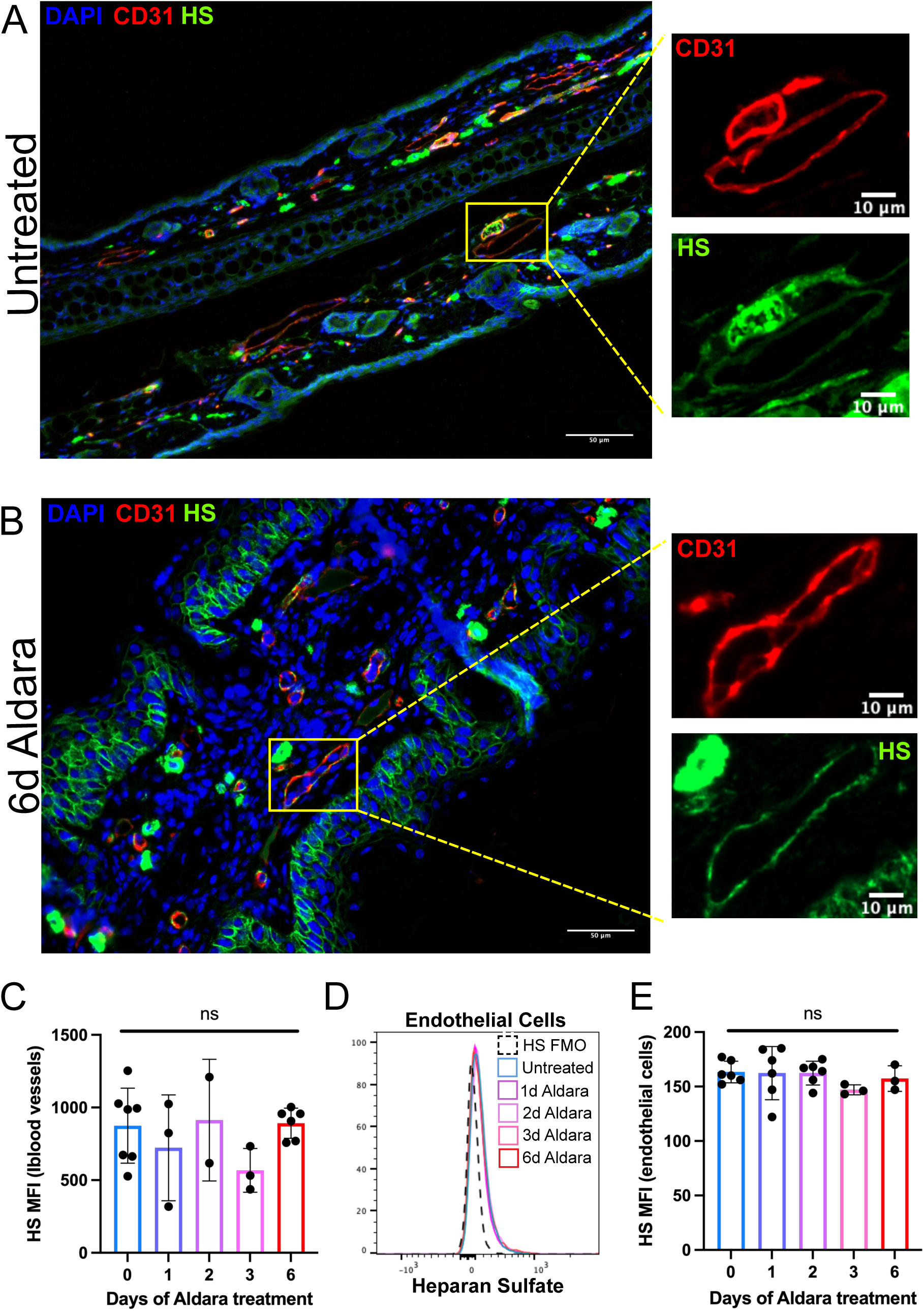
Heparan sulfate in the endothelial glycocalyx is unchanged in response to psoriasis-like skin inflammation. Mice were treated with daily topical application of Aldara cream for 0, 1, 2, 3 or 6 days on both ear pinnae. Skin was stained for HS (green), CD31 (red) and DAPI (blue) (**A, B**) and mean MFI of HS staining measured within the vessel using imageJ (**C**). HS expression on endothelial cells (CD45-CD31+) was measured using flow cytometry (**D**) by calculating the geometric mean fluorescence intensity (gMFI) of HS on CD45-CD31+ cells (**E**). Scale bars in the large images in A and B represent 50 μm, and in the small images 10 μm. **C** and **E** show pooled data from 2 experiments. **A and B** are representative images of skin sections, with zoomed in images of individual vessels highlighted by yellow squares. **D** is a histogram representative of 2 experiments. Data were analysed by one-way ANOVA with Tukey’s multiple comparison test. ‘ns’ denotes not significant. Error bars represent mean ± SD.

### Leukocytes have a cell surface glycocalyx, from which heparan sulfate is lost in response to psoriasis-like skin inflammation

Previous research has suggested several immune subsets including monocytes^20^, neutrophils^21^ and macrophages^22^, may express heparan sulfate on their surface, yet little is known about the function of the leukocyte glycocalyx during inflammation *in vivo*. To determine if the source of circulating HS in inflammation could be immune cells, flow cytometry was used to reveal a clear population of HS+ CD45+ cells in skin (**Fig 3A**), which is reduced in response to psoriasis-like skin inflammation (**Fig 3B**). Uniform Manifold Approximation and Projection (UMAP) visualisation of leukocytes from naïve skin shows widespread expression of HS on the surface of macrophages, monocytes, dendritic cells, T cells, regulatory T cells, and neutrophils (**Fig 3C, D**). Monocytes and neutrophils had the greatest proportion of HS+ cells, as well as the greatest decrease in HS positivity in response to psoriasis-like skin inflammation (**Fig 3E**). A relatively high proportion of regulatory T cells express HS on their surface, however, this was not significantly downregulated under inflammatory conditions (**Fig 3E**).

**Figure 3.**
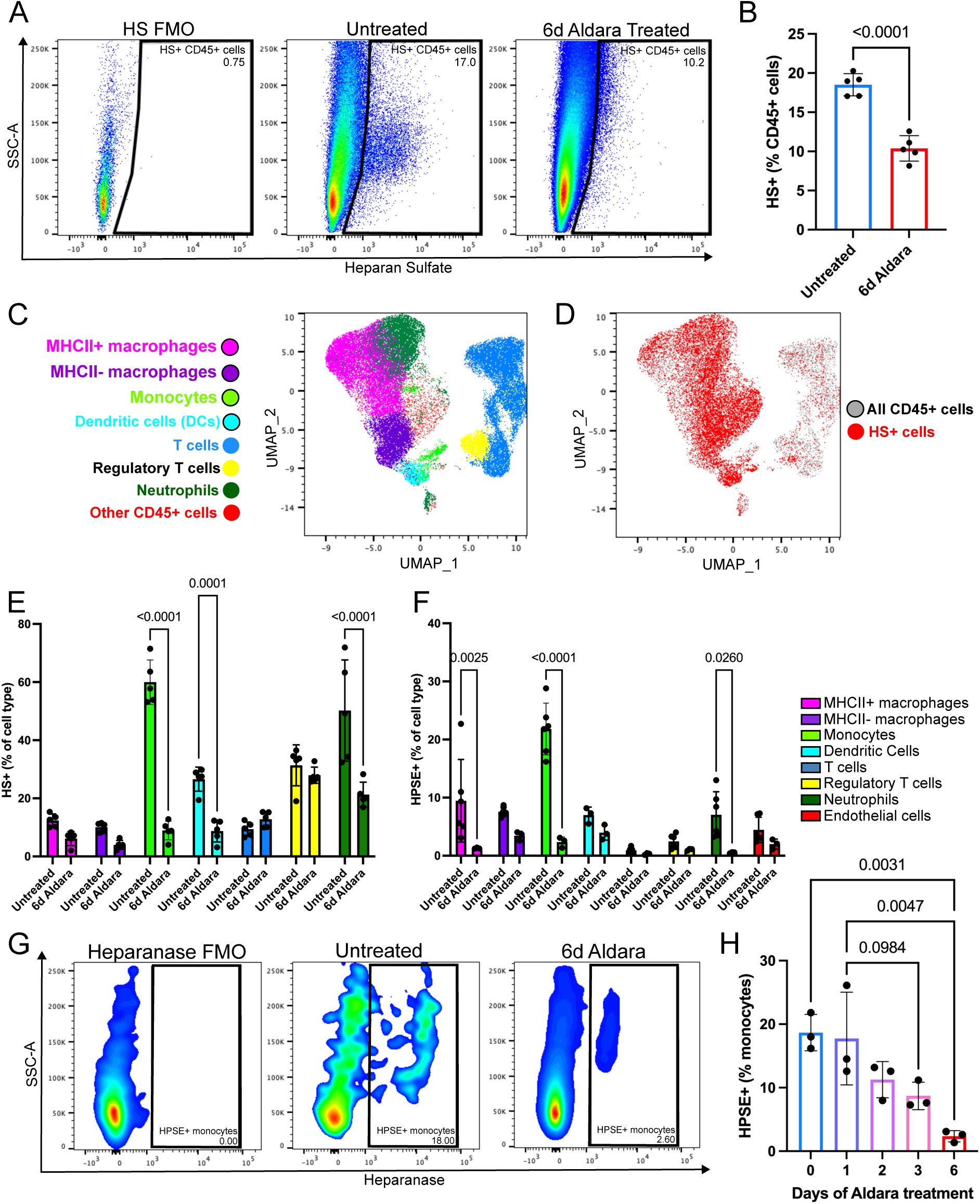
Immune cells lose cell surface heparan sulfate in response to psoriasis-like skin inflammation. Mice were treated topically with Aldara cream to the ear pinnae for 6 days. HS expression on immune cells was measured using flow cytometry (**A**) and the percentage of total CD45+ cells expressing HS was quantified in both untreated and Aldara treated skin (**B**). Naïve mouse skin was stained for flow cytometric analysis and major immune cell types were clustered using the UMAP algorithm (**C**). Cells positive for heparan sulfate (HS) were mapped onto the resulting clusters (**D**). The percentage of leukocyte subsets expressing HS with and without Aldara cream treatment is shown in **E**. Leukocytes were also stained intracellularly for the enzyme heparanase, and the proportions of cells positive for heparanase with and without Aldara cream treatment are shown in **F**. Representative flow plots of monocyte heparanase staining are shown in **G** and the proportions of monocytes positive for heparanase with subsequent days of Aldara cream treatment is shown in **H**. Data show 2 independent experiments, except from **H** which is representative of 2 independent experiments. **A and G** show representative flow plots. Data in **B** were analysed by an unpaired t test, in **E and F** by 2-way ANOVA and **H** by 1-way ANOVA, both with Tukey’s multiple comparisons test. Data with no p values shown are not significant (p>0.05). Error bars represent mean ± SD.

The only mammalian protein capable of cleaving HS along the polysaccharide chain is the endo-glucuronidase, heparanase^28^. Flow cytometric staining for heparanase stored intracellularly in leukocytes revealed that cells of the myeloid lineage had the highest proportion of heparanase positive cells (**Fig 3F**). In particular, a proportion of monocytes show clear heparanase staining (**Fig 3G**) which was reduced with Aldara cream treatment of the skin (**Fig 3H**), possibly indicating its release from the cell. These data suggest that leukocytes have a glycocalyx on their cell surface, which is shed in response to leukocyte-derived heparanase during skin inflammation.

### The heparan sulfate mimetic Tet-29 lessens heparan sulfate shedding on leukocytes and reduces their accumulation in skin during psoriasis-like inflammation

To determine the importance of heparan sulfate loss from the surface of leukocytes, a heparan sulfate mimetic Tet-29 was injected alongside Aldara cream treatment (**Fig 4A**). Zubkova et al.^29^ show that Tet-29 is a potent inhibitor and reduces the cleavage of a heparan sulfate pentasaccharide. Here, flow cytometry revealed that Tet-29 partially protected HS on leukocytes during skin inflammation, resulting in a similar proportion of CD45+ cells staining positively for HS in inflamed Tet-29 treated mice as in naïve mice (**Fig 4B**). Furthermore, Tet-29 treatment resulted in reduced circulating HS in the blood during inflammation compared to PBS treated control mice (**Fig 4C**), indicative of reduced glycocalyx shedding.

**Figure 4.**
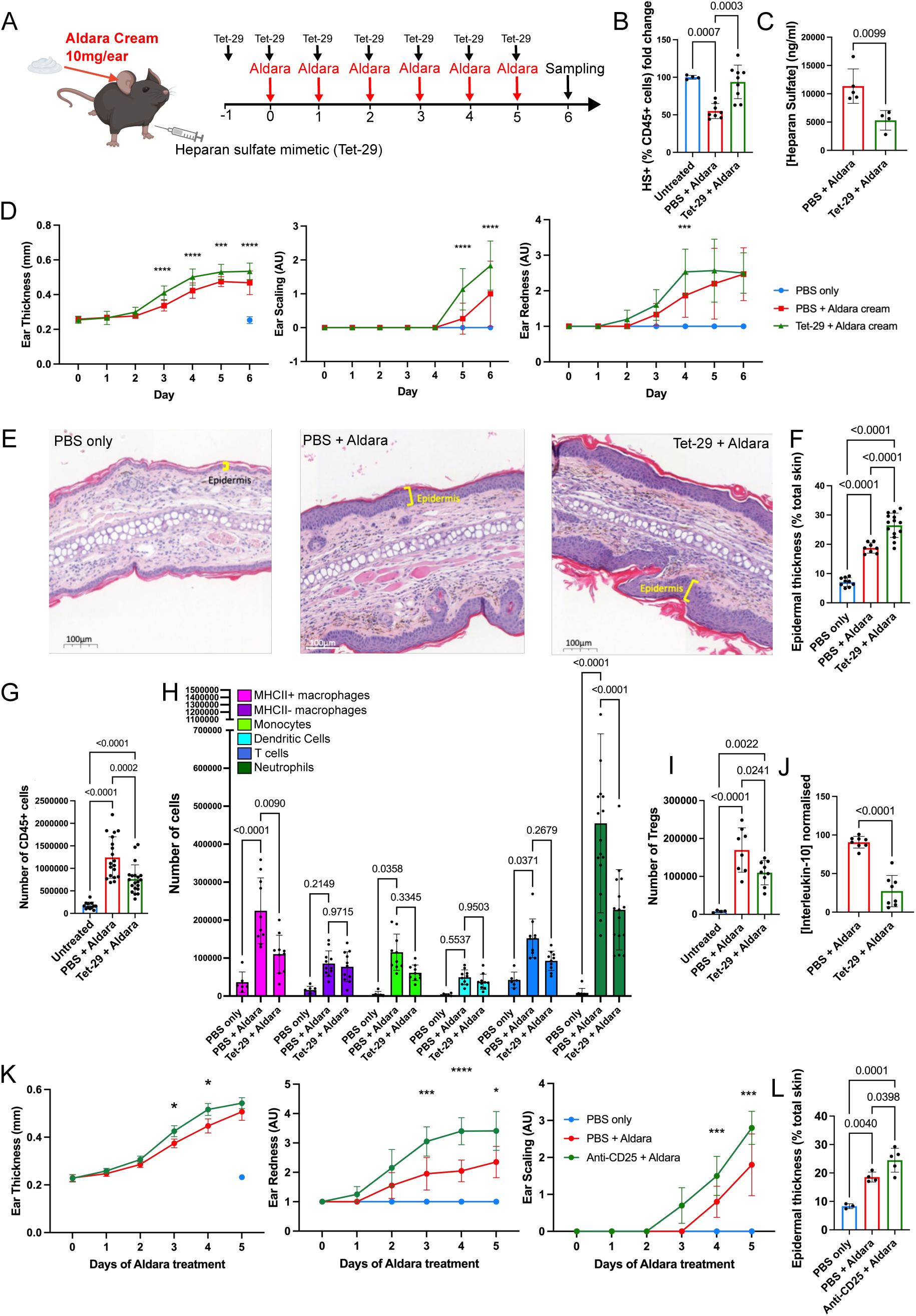
A heparan sulfate mimetic reduces heparan sulfate loss in leukocytes in response to skin inflammation, inhibiting their accumulation in skin, yet enhances clinical signs of inflammation. Mice were treated topically with 10mg Aldara cream to the ear pinnae daily for 6 days (d0 to 5), alongside intraperitoneal injection of the heparan sulfate mimetic, Tet-29, daily for 7 days (d -1 to 5) (**A**), or PBS was injected as a control. Total CD45+ cells were stained for HS and analysed by flow cytometry (**B**). Levels of heparan sulfate in the blood were measured by ELISA (**C**). Clinical signs of inflammation were monitored by measuring ear thickness using callipers and visually scoring ear redness and ear scaling (**D**). Skin was sectioned and stained using haematoxylin and eosin (H&E) (**E**) and epidermal thickness measured from images and calculated as a % of total ear thickness (**F**). Total numbers of CD45+ cells were quantified (**G**) as well as numbers of macrophages, monocytes, dendritic cells, T cells, neutrophils (**H**) and regulatory T cells (**I**) by flow cytometry. Levels of interleukin-10 were measured in skin by ELISA (**J**). Mice were treated with a Treg depleting antibody (anti-CD25) during the course of Aldara cream treatment; ear thickness was measured using callipers and ears were visually scored for redness and scaling (**K**). Aldara and anti-CD25 treated skin was also stained using H&E and epidermal thickness calculated as a % of total ear thickness (**L**). Histology scale bars in **E** show 100 µm. **D, F, G and H** are pooled data from 4 independent experiments. Data in **B, C, I, J, K and L** are pooled from 2 independent experiments. Data in **D, H and K** were analysed using a two-way ANOVA with Tukey’s multiple comparisons test, except for redness and scaling measurements which are discontinuous variables and so a Kruskal-Wallis with Dunn’s multiple comparisons was used. **B, F, G, I and L** were analysed by two-way ANOVA with Tukey’s multiple comparisons test, and **C and J** by unpaired t test. AU = arbitrary units. Error bars represent mean ± SD.

We hypothesised that protecting leukocytes from HS degradation would reduce their migration into skin and as, a result, would lead to reduced psoriasis-like skin inflammation. However, it was found that clinical readouts of inflammation were significantly enhanced with Tet-29 administration alongside Aldara cream treatment when compared to mice injected with phosphate-buffered saline (PBS) as a control (**Fig 4D**). Epidermal thickness, a hallmark of psoriasis severity^30^, was measured using haematoxylin and eosin (H&E) staining (**Fig 4E**) and was found to be significantly greater in inflamed Tet-29 treated mice compared to inflamed control mice (**Fig 4F**). To investigate why inflammation was increased rather than reduced by Tet-29, flow cytometry was used to examine the number of leukocytes in the skin. Counterintuitively, it was found that overall numbers of CD45+ cells in inflamed Tet-29 treated skin were decreased compared to inflamed PBS treated skin, but were higher than in uninflamed skin (**Fig 4G**), suggesting that Tet-29 partially blocks the accumulation of immune cells in skin in response to inflammation. MHCII+ macrophages and neutrophils, were significantly reduced in number in response to Tet-29 treatment (**Fig 4H**). Similar effects were seen in inflamed mice treated with a related HS mimetic^31^, which also reduced leukocyte recruitment into skin and enhanced inflammation (**Fig S3**). These data suggest that protection of HS on the immune cell surface during psoriasis-like skin inflammation reduces the ability of leukocytes to transmigrate into skin.

To investigate why skin inflammation is exacerbated whilst immune cell numbers are reduced in skin in response to Tet-29, the number of skin regulatory T cells (Tregs) was examined. Tregs are an abundant anti-inflammatory cell type in skin, capable of releasing factors such as interleukin-10 (IL-10) to reduce inflammation^32^. Tregs were significantly reduced in number during skin inflammation in Tet-29 treated mice, relative to control mice (**Fig 4I**), suggesting that the heparan sulfate mimetic may reduce the ability of Tregs to enter skin and carry out immunosuppressive functions. Indeed, by ELISA reduced levels of IL-10 were found in Tet-29 treated compared to PBS treated inflamed skin (**Fig 4J**).

To confirm that reduced Treg accumulation leads to enhanced inflammation in Tet-29 treated inflamed mouse skin, mice were treated with Aldara cream alongside injection with anti-CD25 to deplete Tregs (**Fig S4A, S4B**). Depletion of Tregs significantly increased signs of inflammation in skin including ear thickness, redness and scaling (**Fig 4K**), epidermal thickening (**Fig S4C, 4L**), and IL-17 production in the ear draining lymph node (**Fig S4D**).

In addition to Treg depletion, Tet-29 treatment reduced myeloid cells including MHCII+ macrophages and neutrophils in skin, as mentioned above (**Fig 4H**). To determine the impact of neutrophil depletion, an anti-Ly6G antibody was used to deplete neutrophils in the skin of mice with psoriasis-like skin inflammation (**Fig S5A, B**). However, no significant differences in inflammation were observed (**Fig S5D-H**), suggesting that neutrophils do not affect the clinical signs of psoriasis-like skin inflammation. Therefore, in Tet-29 treatment, reduced skin neutrophils will likely not impact the severity of inflammation.

iCCR KO mice, lacking CCRs 1, 2, 3 and 5^33^ have reduced numbers of myeloid cells in skin following treatment with Aldara cream, including DCs, monocytes and some macrophages (**Fig S6B)**. Despite this, the iCCR KO mice showed no differences in psoriasis-like skin inflammation severity compared to littermate controls (**Fig S6E-I**). These findings demonstrate that the recruitment of myeloid cells to the skin does not affect the severity of psoriasis-like skin inflammation and therefore in Tet-29 treated mice, the reduced myeloid cells are unlikely to affect the severity of inflammation.

These data indicate that reduced HS shedding from the leukocyte surface in response to psoriasis-like skin inflammation decreases immune cell accumulation in the skin. This exacerbates skin inflammation due to reduced skin Tregs and the associated reduction in anti-inflammatory cytokines.

### Loss of cell surface heparan sulfate results in reduced monocyte migration *in vitro*

To determine whether the loss of cell surface HS on leukocytes during inflammation promotes migration into inflamed tissues, heparan sulfate was enzymatically degraded on the surface of cells from murine bone marrow using bacterial heparinase, and the ability of these cells to migrate towards CCL7 was investigated (**Fig 5A**). Flow staining confirmed the successful degradation of HS on the surface of bone marrow monocytes treated with heparinase (**Fig 5B-C**), and showed that heparinase treatment promotes the migration of monocytes towards chemokine (**Fig 5D**). These data show that leukocytes have an increased ability to migrate when they have reduced HS on their cell surface, providing a potential explanation for how the heparan sulfate mimetic Tet-29 may be responsible for reduced numbers of immune cells in the skin during inflammation. This suggests that the reduction in leukocyte HS under inflammatory conditions may *in vivo* be a mechanism to aid the recruitment of immune cells to the site of inflammation.

**Figure 5.**
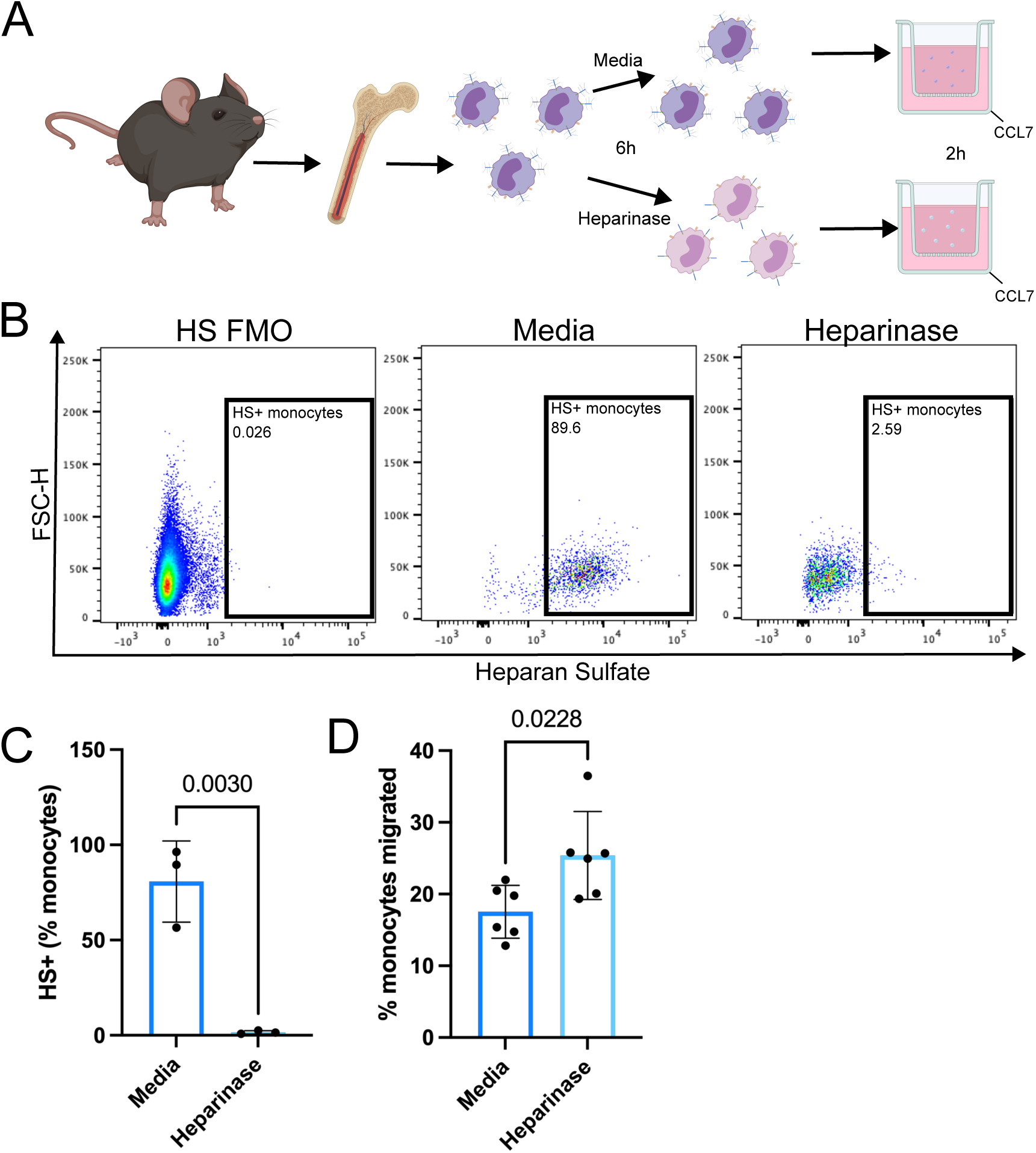
Degradation of heparan sulfate on the surface of monocytes increases their ability to migrate in vitro. Mouse bone marrow was cultured with or without heparinase I and III before being added to the top well of a transwell (**A**). Cells were allowed to migrate towards CCL7 for 2h and migrated cells were collected stained for heparan sulfate and analysed by flow cytometry (**B**) and the proportion of transmigrated monocytes expressing HS was calculated (**C**) along with the percentage of total monocytes that had migrated (**D**). Data are from 2 independent experiments each with 3 technical replicates. **B** shows representative flow plots. Data were analysed using an unpaired t test. Error bars represent mean ± SD.

## DISCUSSION

The remodelling of the glycocalyx on the endothelium and its role in facilitating leukocyte recruitment has been increasingly studied over the last 20 years, particularly in the context of inflammatory diseases^34^. Yet, little is known about the function of its counterpart; the leukocyte surface glycocalyx, and how it may be altered during inflammation. In the present study, we have demonstrated the presence of a leukocyte glycocalyx *in vivo* and shown its remodelling in response to skin inflammation, suggesting a novel mechanism for *in vivo* leukocyte recruitment into skin.

Whilst the breadth of glycocalyx research has expanded in recent years, it remains primarily focussed on the endothelial glycocalyx. Li et al^19^ demonstrate that human patients with psoriasis have lower levels of HS on the dermal vascular endothelium than in healthy patients, yet the leukocyte glycocalyces in these patients were not examined. This common focus on the endothelial glycocalyx has produced assumptions. For example, increased levels of circulating glycocalyx components, such as HS and protein cores, are assumed to be indicative of their shed from the endothelial glycocalyx. Our findings of a glycocalyx on the immune cell surface challenge this assumption.

We propose that the loss of HS from the leukocyte surface may contribute to the elevated circulating HS found in inflamed mice in our study. Although, it is likely that other cell types contribute to this, including fibroblasts and keratinocytes, both of which produce HS^35^. These data emphasise the importance of more work on the cell-specific function of HS across a variety of cell types.

Much of the current literature on heparanase-mediated degradation of the endothelial glycocalyx has focussed on endothelial cells as the primary source of heparanase. Here it was shown that myeloid cells are another important source of the enzyme, a finding supported by Arokiasamy et al^36^ who identify macrophages as a key source of heparanase. This suggests a more widespread expression of heparanase and context dependent effects.

Having demonstrated the presence of an HS-containing glycocalyx on immune cells, we next examined its function. Enzymatic degradation of leukocyte HS enhanced the cells’ ability to migrate through a transwell. Sabri et al^37^ showed a similar finding, with THP1 monocytes shedding a proportion of their cell surface glycocalyx in response to IFNγ treatment, resulting in increased cell adhesion to antibody-coated spheres. Shedding the leukocyte glycocalyx may result in increased exposure of chemokine receptors on the cell surface, allowing monocytes to more readily bind chemokines, resulting in enhanced migration (**Fig 6**). *In vivo*, leukocyte glycocalyx shedding may also expose cell surface adhesion molecules, allowing for greater adhesion to endothelial cells and subsequent transmigration. However, other *in vitro* studies^38, 39^ show opposite effects of HS in migration, with reduced glycosaminoglycans on the cell surface associated with their reduced ability to bind chemokine. These data suggest the functions of leukocyte glycocalyx shedding may be context dependent and complex beyond simply shielding and exposing adhesion molecules.

**Figure 6.**
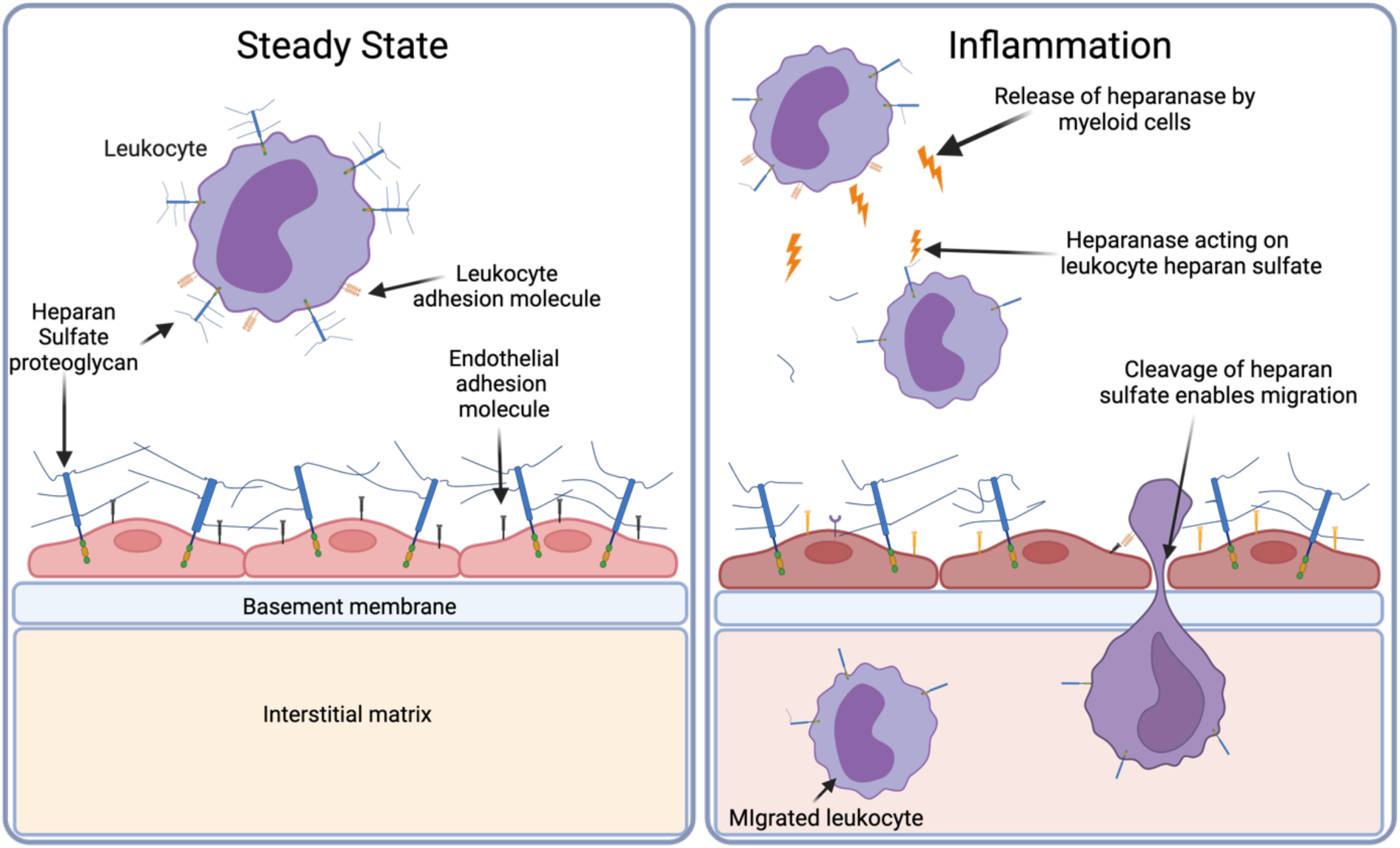
Heparan sulfate shedding from the leukocyte glycocalyx enables their infiltration into the skin during psoriasis-like skin inflammation. We propose that the leukocyte glycocalyx acts as a barrier to migration and is shed during inflammation to enable migration to inflammatory sites. Adapted from Sutherland et al. 2023.

Over recent years, heparanase inhibitors have been developed to treat inflammatory diseases^40–42^ and cancers^43–46^. Our data supports the idea that the HS mimetic, Tet-29, inhibits heparanase *in vivo*. Peck et al.^47^ treated mice with Tet-29 during a model of multiple sclerosis which resulted in reduced leukocyte recruitment in the central nervous system. Here, we expand on these findings by providing a potential mechanism for how it may do so, via the protection of HS on the leukocyte surface. However, it is important to note that Tet-29 may also affect other components of the migration process such as binding to cytokines^48^, chemokines^11^ or integrins^49^.

Our findings demonstrate that heparanase inhibitors have complex effects on the immune system in this context. We hypothesise that in our model, pro-inflammatory cell recruitment is not the sole driver of inflammation, as γδ T cells and group 3 innate lymphoid cells which drive the inflammation^50^ are largely skin resident^51^, yet Tregs are more dependent on recruitment to the skin^52^. Therefore, reducing immune cell recruitment to the skin using Tet-29 in this model only marginally affects pro-inflammatory cell activity, but more staunchly affects anti-inflammatory Treg migration, resulting in the enhanced inflammation observed here.

Based on our findings, we propose that the recruitment of monocytes into skin during psoriasis-like skin inflammation and their release of heparanase results in the cleavage of HS, allowing subsequent immune cells to migrate into skin more readily (**Fig 6**), a finding supported by previous studies^53–56^. Furthermore, monocytes may be able to cleave HS on their own cell surfaces, as well as on the surface of other immune cells, in order to facilitate their recruitment to inflamed skin. Together, these findings demonstrate the importance of HS proteoglycans in regulating leukocyte recruitment in inflammatory disease. However, they challenge the model that these proteoglycans are primarily endothelial and emphasise the importance of HS on leukocytes themselves.

## MATERIALS AND METHODS

### Mice

All animal experiments were locally ethically approved and performed in accordance with the UK Home Office Animals (Scientific Procedures) Act 1986. C57BL/6 mice were obtained from Charles River Laboratories. ICCR KO mice were created by Professor Gerard Graham (University of Glasgow, UK) on a C57BL/6N background^33^ and were bred and maintained in specific pathogen-free conditions in house. Mice were 7-14 weeks old and female, except for ICCR mice which were mixed sex.

### Induction of psoriasis-like skin inflammation

Psoriasis-like skin inflammation was induced as previously described in Linley et al^24^. Briefly, mice were anaesthetised and treated daily with 10 mg of Aldara cream (Meda Pharmaceuticals) containing 5% Imiquimod, by topical application on both ear pinnae for 6 days (days 0 to 5). Ear thickness was measured daily using a digital micrometer (Mitutoyo) and ear scaling and redness scored daily.

### Glycocalyx and Immune cell modulation

#### Heparanase inhibition

To inhibit degradation of heparan sulfate, mice were injected intraperitoneally with 600 μg of the heparan sulfate mimetic Tet-29^28^, kindly provided by Dr Olga Zubkova (University of Wellington, New Zealand), or with phosphate buffered saline as a control, daily for 7 days (days -1 to 5), alongside Aldara cream treatment (given on days 0 to 5, as described above).

#### Regulatory T cell depletion

Mice were injected intraperitoneally with 250 μg anti-CD25 antibody (BioXCell, clone PC-61.5.3) or with phosphate buffered saline as a control, on days -2, 0, 2 and 4, alongside Aldara cream treatment (given on days 0 to 5, as described above).

#### Neutrophil depletion

Mice were injected intraperitoneally with 500 μg anti-Ly6G antibody (BioXCell, clone 1A8) or with 500 μg isotype control (BioXCell), on days -2, 0, 2 and 4, alongside Aldara cream treatment (given on days 0 to 5, as described above).

### Haematoxylin and eosin staining

Ear tissue was fixed in 10% neutral buffered formalin (Sigma Aldrich), then embedded in paraffin and cut to 5 μm. Haematoxylin and eosin staining was carried out using an automated Shandon Varistain V24-4. Images were acquired using a 3D HISTECH Pannoramic-250 microscope slide-scanner (3D HISTECH). Snapshots and measurements were taken with Case Viewer software (3D HISTECH). Thickness measurements of both the epidermis and of the whole ear were taken at 3 different points on each section, and epidermal thickness was quantified as a percentage of total skin thickness. At least 3 sections were measured per mouse and data points represent the mean epidermal thickness per mouse.

### Cell isolation

Ears were removed from euthanised mice and were split in half. Tissue was digested with 0.5 mg/ml DNAse I (Roche) and 0.25 mg/ml Liberase TM (Roche) in complete RPMI (RPMI-1640 (Sigma-Aldrich) supplemented with 10% heat-inactivated FBS (Life Technologies Ltd), 1% Penicillin-Streptomycin solution, 1 mM Sodium Pyruvate, 2 mM L-glutamine, 25 nM 2-β-mercaptoethanol, 1x non-essential amino acid solution, 20 mM HEPES buffer; (all Sigma-Aldrich)), shaking for 2 hours at 37⁰C before disaggregation in a Medimachine System (BD BioSciences) for 6 minutes. Debris was removed by filtering through a 70 µm cell strainer before cells were washed and counted using Trypan blue exclusion and a haemocytometer.

Ear-draining (auricular) lymph nodes (LN) were collected from mice in complete RPMI and single cell suspensions were acquired by gentle agitation on a 70μm strainer.

Blood samples were collected via cardiac puncture under terminal anaesthesia and were placed directly into in EDTA (Lonza). Red blood cells were lysed using ammonium-chloride-potassium (ACK) lysis buffer (Gibco).

Bone marrow cells were collected by centrifugation of fibulas and tibias, then red blood cells were lysed using ACK lysis buffer (Gibco).

### Flow Cytometric analysis

Cells examined for cytokine production were resuspended in a stimulation media made of complete RPMI (see above) plus 10 µM Brefeldin A, 50 ng/mL phorbol 12-myristate 13-acetate (PMA) and 500 ng/mL ionomycin (all Sigma Aldrich) and incubated for 4 hours at 37 °C. Cells were incubated with a Live/dead blue fixable stain (Invitrogen, ThermoFisher Scientific) diluted to 1:2000 with PBS and Fc block (see **Table S1**) then stained with extracellular antibodies (**Table S1**) in FACS wash (PBS with 1% FBS and 2mM EDTA). Cells were fixed using Foxp3/transcription factor staining buffer set (eBioscience). Where indicated, overnight staining using a biotinylated anti-heparan sulfate antibody (# in **Table S1**) followed by a secondary fluorescently labelled streptavidin antibody (**Table S1**) was performed. Where intracellular staining was performed, cells were permeabilised using the Foxp3/transcription factor staining buffer set (eBioscience) and cells were incubated for 2h in intracellular antibodies (* in **Table S1**) in permeabilisation buffer (eBioscience). Where appropriate, cells were also stained with an anti-rabbit IgG to detect the anti-heparanase antibody (**Table S1**). Data were acquired with a Fortessa X20 (Becton Dickinson) flow cytometer using BD FACSDiva Software. Data were analysed using FlowJo (version 10.8.0) (Treestar Inc.) Gating on cells was carried out according to the gating strategy in **Fig S2**.

### Uniform manifold approximation and projection (UMAP) clustering

UMAP plots were generated using R^57^ by clustering cells based on expression of the markers CD11b, CD11c, CD3, F4/80, Foxp3, Ly6C, Ly6G, MHCII, TCRβ and TCRγδ.

### Enzyme-linked immunosorbent assay (ELISA)

Skin was snap frozen before bead beating using a TissueLyser II (Qiagen) at 20 Hz for 2 minutes with a 5mm ball bearing. Tissue was then placed in lysis buffer (0.1% Triton X100 (Sigma-Aldrich) with cOmplete mini protease inhibitors (Roche) in phosphate buffered saline) for 30 minutes at 4 °C.

Blood samples were taken from mice by cardiac puncture under terminal anaesthesia and left on ice to coagulate for 4 hours. Blood was centrifuged at 15,000 x g for 10 mins at 4 °C and supernatants collected and frozen.

Levels of heparan sulfate and interleukin-10 were measured using enzyme-linked immunosorbent assay kits from Finetest and R&D systems respectively. Data depicts the mean of duplicates for each sample.

### Immunofluorescence staining

Sections were dewaxed in xylene and rehydrated in decreasing concentrations of ethanol. Antigen retrieval was carried using Tris-EDTA buffer (pH 9.0) before permeabilization in 0.5 % Triton-X100 (Sigma-Aldrich). Sections were blocked using donkey serum in 1% BSA for 30 mins at room temperature, then using an Avidin/Biotin blocking kit (Vector) according to manufacturer’s instructions. Sections were further blocked using Mouse Ig Blocking Reagent (Mouse on Mouse Immunodetection Kit, Vector) for 1 hour. Sections were incubated with primary antibodies (**Table S2**) overnight at 4 degrees, followed by anti-heparan sulfate antibody (**Table S2**) for 10 minutes, followed by tertiary antibodies against HS and secondary antibodies against CD31 (**Table S2**) for 1 hour at room temperature, all made up in MOM diluent (Mouse on Mouse Immunodetection Kit, Vector). Nuclei were visualised using DAPI (49,6-diamidino-2-phenylindole) (Thermofisher) staining (0.2 µg/ml for 5 minutes) and sections were mounted using Prolong Gold antifade mountant (Thermofisher). Images were collected on a Zeiss Axioimager.D2 upright microscope using 10x, 20x, 40x and 63x EC Plan-neofluar objective lenses and captured using a Coolsnap HQ2 camera (Photometrics) through Micromanager software v1.4.23. Images were processed and analysed using ImageJ (version 1.53a).

### Immunofluorescence image analysis

HS expression was quantified in images by selecting endothelial cells (using CD31) on the perimeter of blood vessels excluding the lumen, and measuring the mean fluorescence value for HS staining within these areas. At least 3 vessels were measured per section and at least 3 sections were measured per mouse. Data represent the MFI (mean fluorescence intensity) per mouse. Vascularisation of skin was quantified by measuring the number of pixels staining positively for CD31 and dividing this by the number of pixels within the boundary of the skin section. At least 5 sections were measured per mouse.

### *In vitro* heparinase treatment

Bone marrow cells were collected by centrifugation of fibulas and tibias, then red blood cells were lysed using ammonium-chloride-potassium (ACK) lysis buffer (Gibco). Cells were resuspended at 2.5 million cells/mL and treated with 1 U/mL heparinase I and III from flavobacterium heparinum (Sigma-Aldrich) for 6 hours at 37°C in complete RPMI.

### Migration experiments

Migration assays were performed using 24-well transwell plates with 5 μm pore size filter inserts (Corning). CCL7 was added to 100 nM in the bottom well in complete RPMI. 100μL bone marrow containing 1.25×10^7^ cells was added to the upper chamber and cells were allowed to migrate for 2 hours at 37 °C, before migrated cells were collected and analysed by flow cytometry. The proportion of cells migrated was calculated by dividing the number of migrated cells by the number of cells recovered from a well with no transwell insert. Data are from 2 independent experiments, performed in triplicate.

### Statistical Analysis

All data were analysed using GraphPad Prism 10. Data were tested for normality using a Shapiro-Wilks test. Unpaired t tests (2 groups) or one-way ANOVAs with Tukey’s multiple comparison test (more than 2 groups) were used for parametric data. Non-parametric data were analysed using Mann-Whitney U tests (2 groups) or Kruskal Wallis with Dunn’s multiple comparisons test (more than 2 groups). Data annotated as ‘fold change’ is normalised as a percentage of the average value of the PBS control group in each experiment.

## Supplementary Materials

**Figure S1.** Characterising changes in clinical signs of inflammation and the vasculature during psoriasis-like skin inflammation.

**Figure S2.** *Flow cytometry gating strategy for leukocytes and endothelial cells*.

**Figure S3.** A related heparan sulfate mimetic inhibits leukocyte recruitment in skin but enhances clinical signs of inflammation.

**Figure S4.** Anti-CD25 treatment depletes regulatory T cells and enhances some signs of inflammation during psoriasis-like skin inflammation.

**Figure S5.** *Depletion of neutrophils does not recapitulate the enhanced inflammation seen with heparanase inhibition during psoriasis-like skin inflammation*.

**Figure S6.** *Depletion of myeloid cells does not recapitulate the enhanced inflammation seen with heparanase inhibition during psoriasis-like skin inflammation*.

**Table S1.** Antibodies used for flow cytometry staining.

**Table S2.** Antibodies for immunofluorescence staining.

## Acknowledgments

The authors acknowledge members of the Saunders lab and Dyer lab for critical discussion. We acknowledge members of the biological services, flow cytometry, histology and bioimaging facilities at the University of Manchester. We acknowledge the gift of ICCR KO mice from Professor Graham from the University of Glasgow. OZ thanks Dr. Andrew Lewis and Dr. Yinrong Lu for an exceptional NMR and mass spectroscopy service, and Ye Li for assistance with the synthesis. Diagrams created using Biorender.com.

## Funding

Sir Henry Dale fellowship jointly funded by the Wellcome Trust and Royal Society 218570/Z/19/Z (DD)

Wellcome Trust center grant 203128/A/16/Z (DD)

Sir Henry Dale fellowship jointly funded by the Wellcome Trust and Royal Society 109375/Z/15/Z (AS)

New Zealand Ministry of Business, Innovation and Employment RTVU1801 (OZ) New Zealand Breast Cancer Foundation R1703 (OZ)

Wellcome Trust Immunomatrix in Complex Disease (ICD) PhD studentship 218491/Z/19/Z (MP)

## Author contributions

Conceptualization: MP, DD, AS

Methodology: MP, IM, AS, DD

Investigation: MP, AH, AS, DD

Visualization: MP

Resources: OZ, SSS

Funding acquisition: DD, AS

Project administration: DD, AS

Supervision: DD, AS

Writing – original draft: MP

Writing – review & editing: MP, DD, AS, SSS

## Data and materials availability

Any data not explicitly evident in the paper will be available on reasonable request.

