## Supplementary information for "Leukocytes have a heparan sulfate glycocalyx that regulates recruitment during inflammation"

### Supplementary material

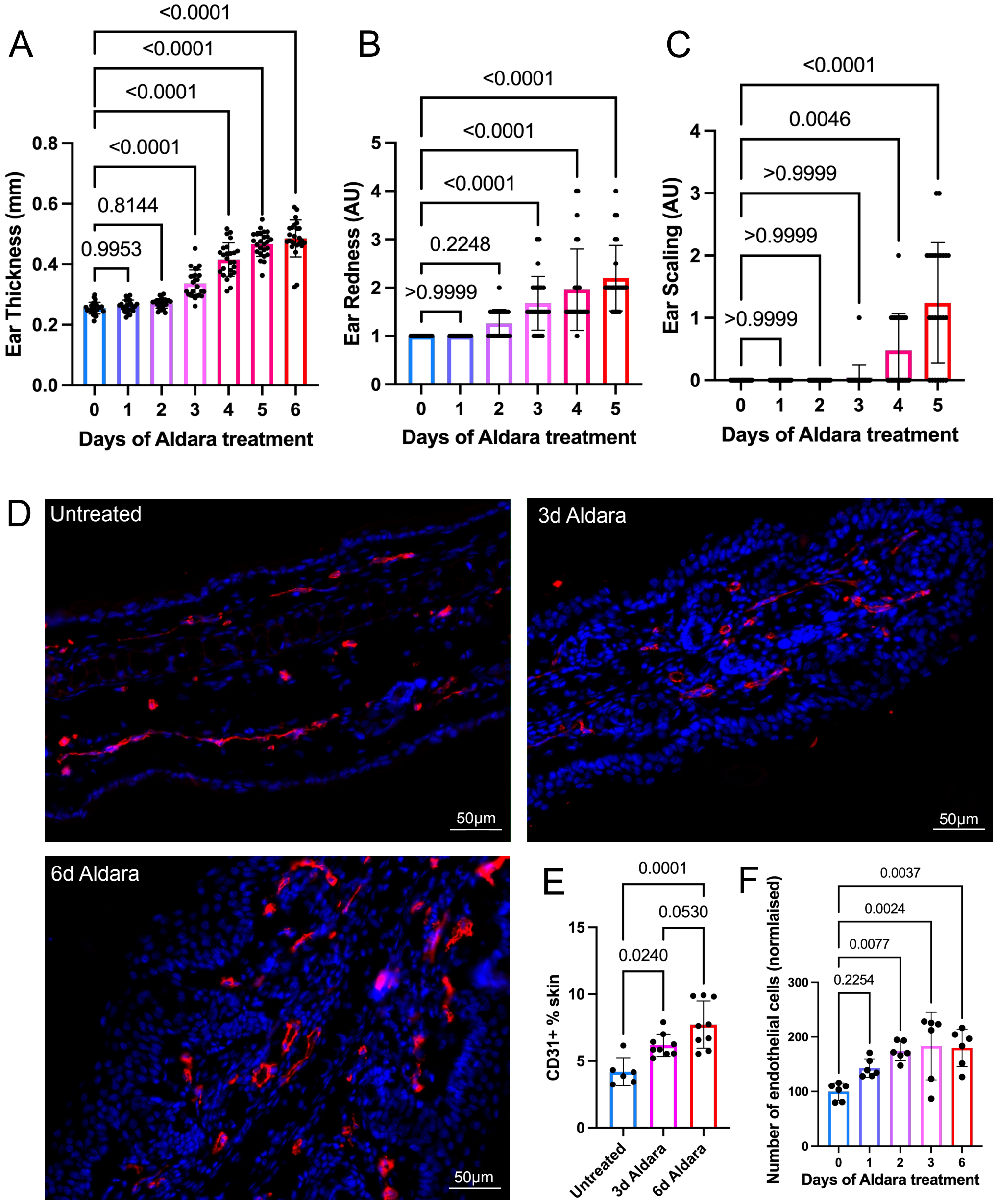


**Figure S1.** *Characterising changes in clinical signs of inflammation and the vasculature during psoriasis-like skin inflammation.* Mice were treated topically with 10mg Aldara cream to each ear pinnae daily for 0-6 days. Clinical signs of inflammation were quantified by measuring ear thickness (**A**) and scoring redness (**B**) and scaling (**C**) against a scale. Skin was sectioned and stained for DAPI (blue) and CD31 (red) as a marker of endothelial cells (**D**) and the vascularisation of the tissue quantified by calculating the % of skin positive for CD31 staining (**E**). The number of endothelial cells was quantified by staining skin for CD31 by flow cytometry and normalising to the mean of untreated controls (**F**). Scale bars in D represent 50µm. A-C show data from 4 pooled experiments. E and F shows data pooled from 2 experiments. D shows representative images from 1 experiment. Data in **A, E and F** were analysed by one-way ANOVA with Tukey's multiple comparison test. Data in **B and C** are discontinuous variables and so were analysed using non-parametric Kruskal-Wallis test with Dunn's multiple comparison test. **A-C** show significant differences compared to untreated mice (day 0) only. Data in **E and F** with no p values shown is not significant (p>0.05). 'AU' denotes arbitrary units. Error bars represent mean ± SD.

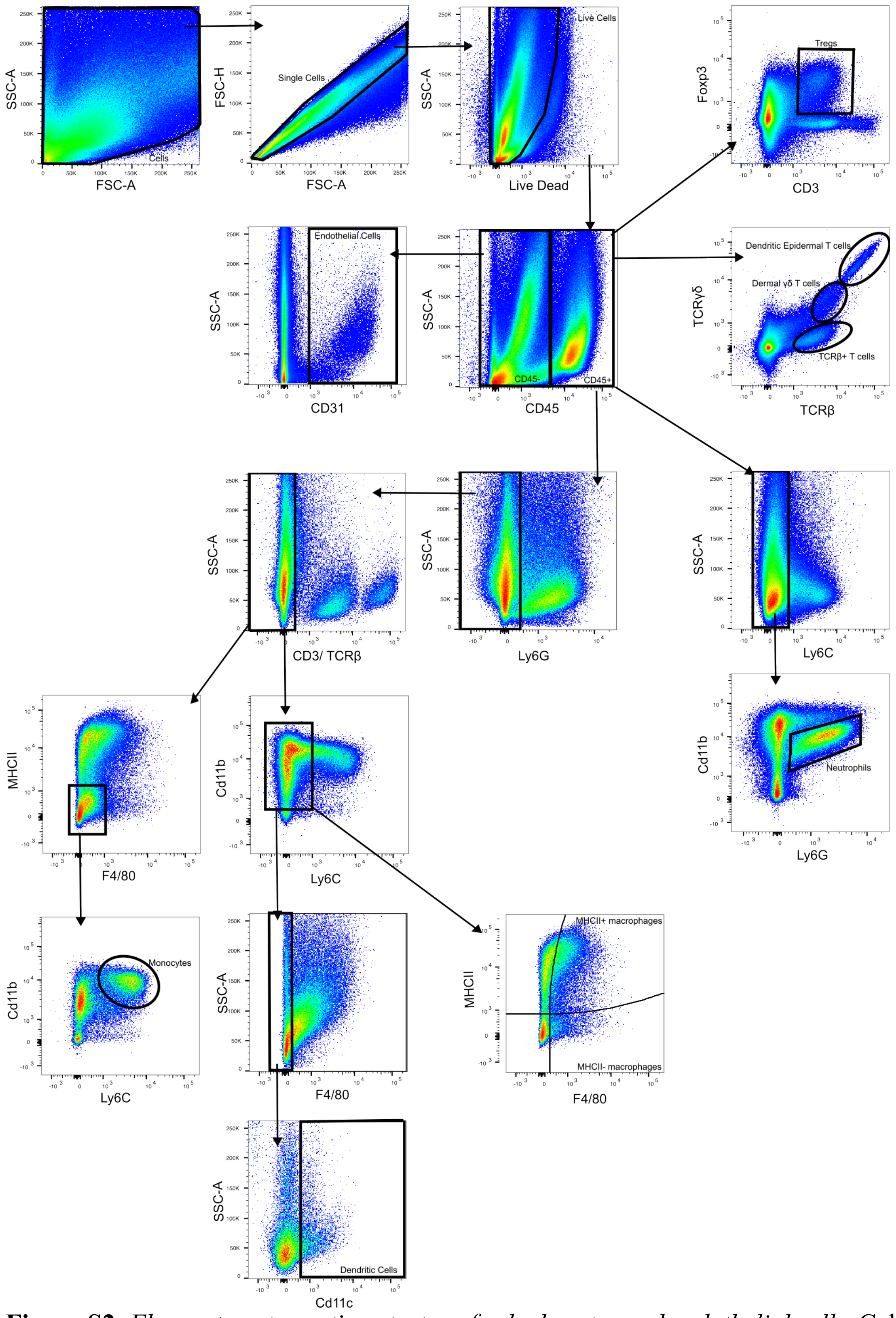


**Figure S2**. *Flow cytometry gating strategy for leukocytes and endothelial cells*. Cells were gated on based on forward scatter (FSC) and side scatter (SSC) characteristics, then single cells were gated on based on linearity of FSC-A versus FSC-H parameters. Live cells were gated on as LiveDead-. Endothelial cells were gated on as CD45-, CD31+. Immune cells were gated on as CD45+ and subsequent gating into subsets is shown.

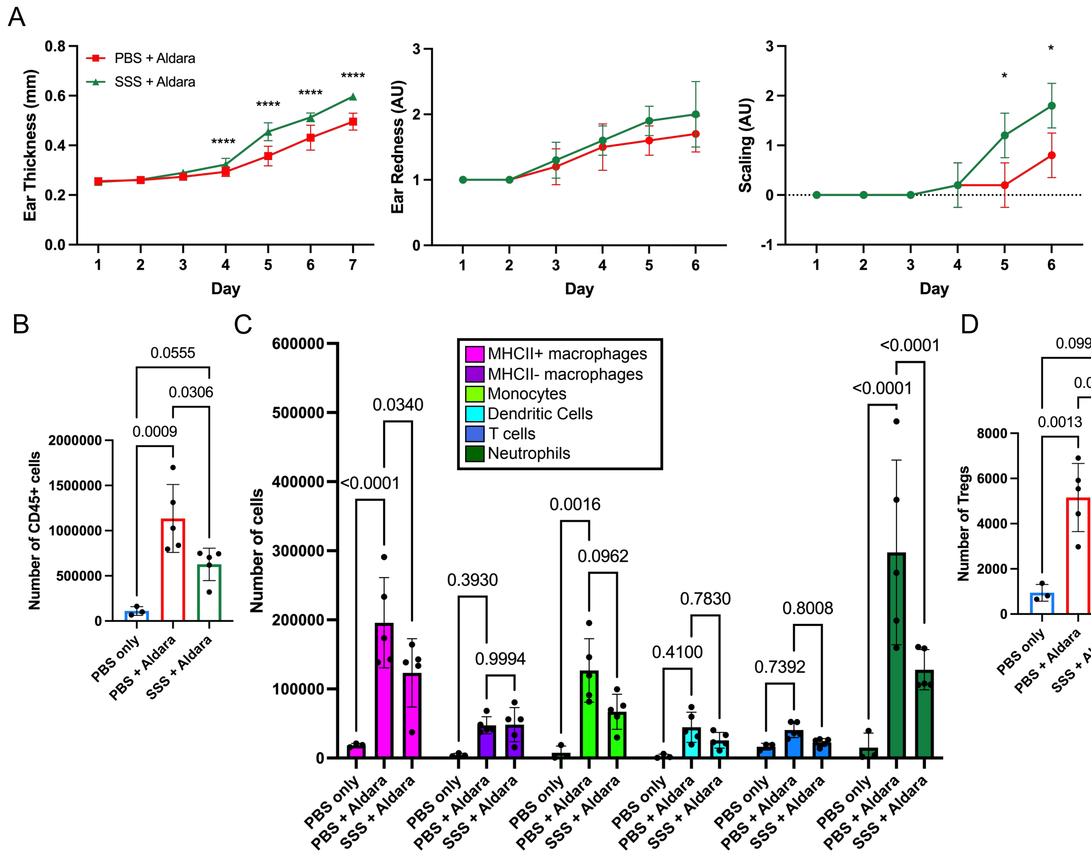


**Figure S3.** *A related heparan sulfate mimetic inhibits leukocyte recruitment in skin but enhances clinical signs of inflammation.* Groups of mice were treated with topical application of 10mg Aldara cream to the ear pinnae daily for 6 days (d0 to 5), alongside intraperitoneal injection of a heparan sulfate mimetic, denoted as ‘SSS’, daily for 7 days (d -1 to 5), or PBS was injected as a control. Clinical signs of inflammation were monitored by measuring ear thickness using callipers and visually scoring ear redness and ear scaling (**A**). Total numbers of CD45+ cells were quantified (**B**) as well as numbers of macrophages, monocytes, dendritic cells, T cells, neutrophils (**C**) and regulatory T cells (**D**). Data are from 1 experiment. Data in **A and C** were analysed using a two-way ANOVA with Tukey’s multiple comparisons test, except for redness and scaling measurements which are discontinuous variables and so a Kruskal-Wallis with Dunn’s multiple comparisons was used. **B, and D** were analysed by two-way ANOVA with Tukey’s multiple comparisons test. AU = arbitrary units. Error bars represent mean ± SD.

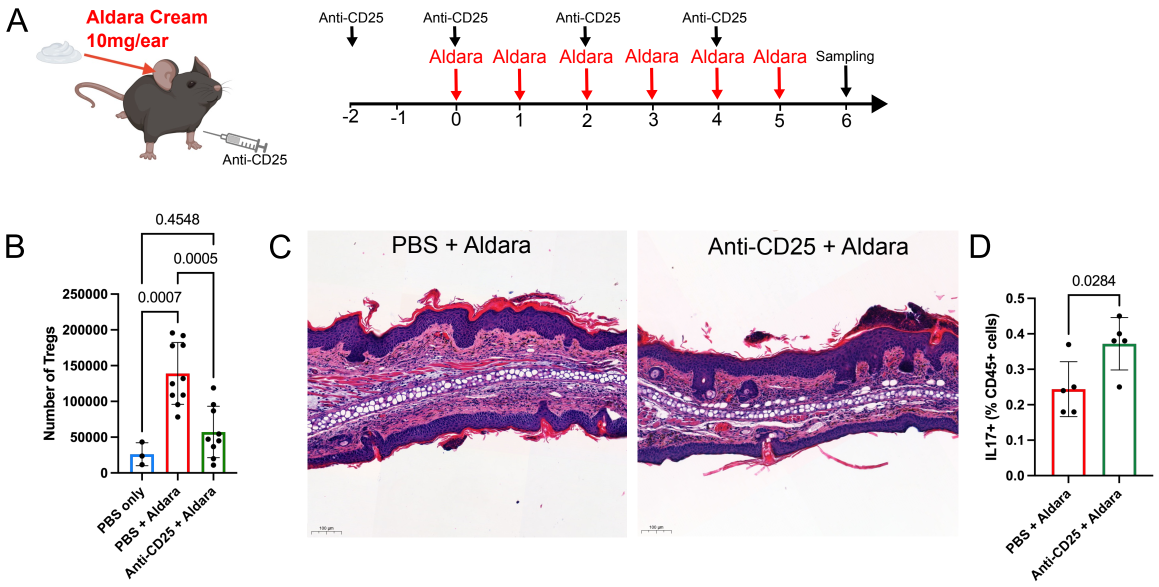


**Figure S4.** *Anti-CD25 treatment depletes regulatory T cells and enhances some signs of inflammation during psoriasis-like skin inflammation.* Mice were treated daily with topical application of Aldara cream to the ear pinnae (d0-6) alongside intraperitoneal injection of an anti-CD25 antibody on days -2, 0, 2 and 4 (**A**), leading to the depletion of Tregs in the skin measured by flow cytometry (**B**). Skin was sectioned and stained using haematoxylin and eosin (H&E) (**C**). Ear draining (auricular) lymph nodes were stained for flow cytometry and the % of CD45+ cells producing interleukin-17 were quantified (**D**). Histology scale bars in **C** represent 100µm. Data shows 2 pooled experiments, except from **C** which shows representative images at 10x magnification. Data were analysed by one-way ANOVA and Tukey's multiple comparison test, except from **D** which was analysed by unpaired t test. Error bars represent mean ± SD.

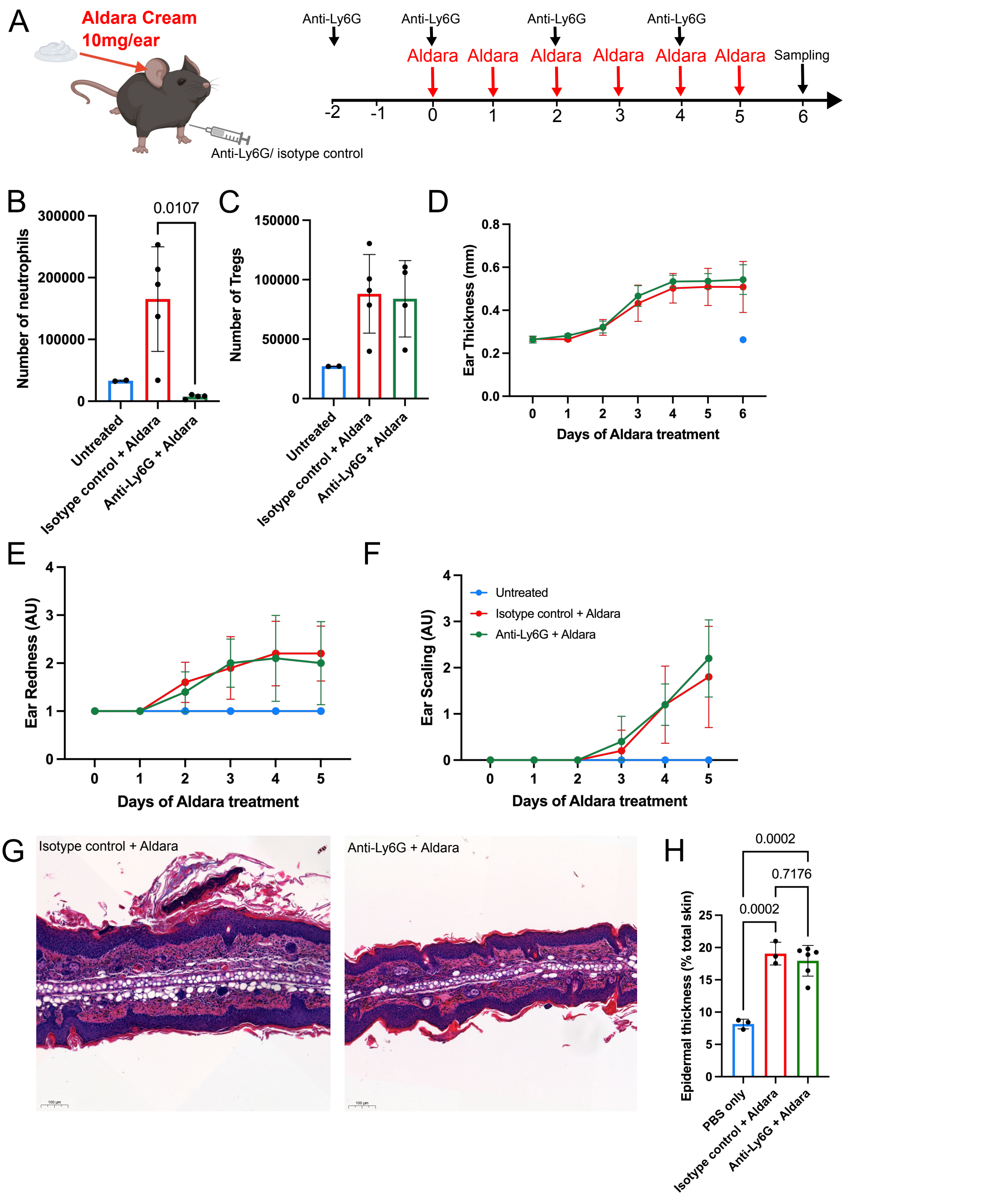


**Figure S5**. *Depletion of neutrophils does not recapitulate the enhanced inflammation seen with heparanase inhibition during psoriasis-like skin inflammation.* Mice were treated daily with topical application of Aldara cream to the ear pinnae (days 0-6) alongside intraperitoneal injection of an anti-Ly6G antibody on days -2, 0, 2 and 4 (**A**), leading to the depletion of neutrophils in the skin measured by flow cytometry (**B**). Treg numbers per 2 ears were also measured by flow cytometry (**C**). Clinical readouts of inflammation were scored including ear thickness (**D**), redness (**E**) and scaling (**F**). Skin was sectioned and stained using haematoxylin and eosin (H&E) (**G**) and epidermal thickness measured and calculated as a % of total ear thickness (**H**). Histology scale bars in **G** represent 100µm. Data shows 1experiment, from which **G** which shows representative images at 10x magnification. Data were analysed by one-way ANOVA and Tukey's multiple comparison test, except from **E**

**and F** which are discontinuous variable and so were analysed by a non-parametric Kruskal-Wallis test with Dunn's multiple comparisons. Data with no p values shown are not significant (p>0.05). Error bars represent mean ± SD.

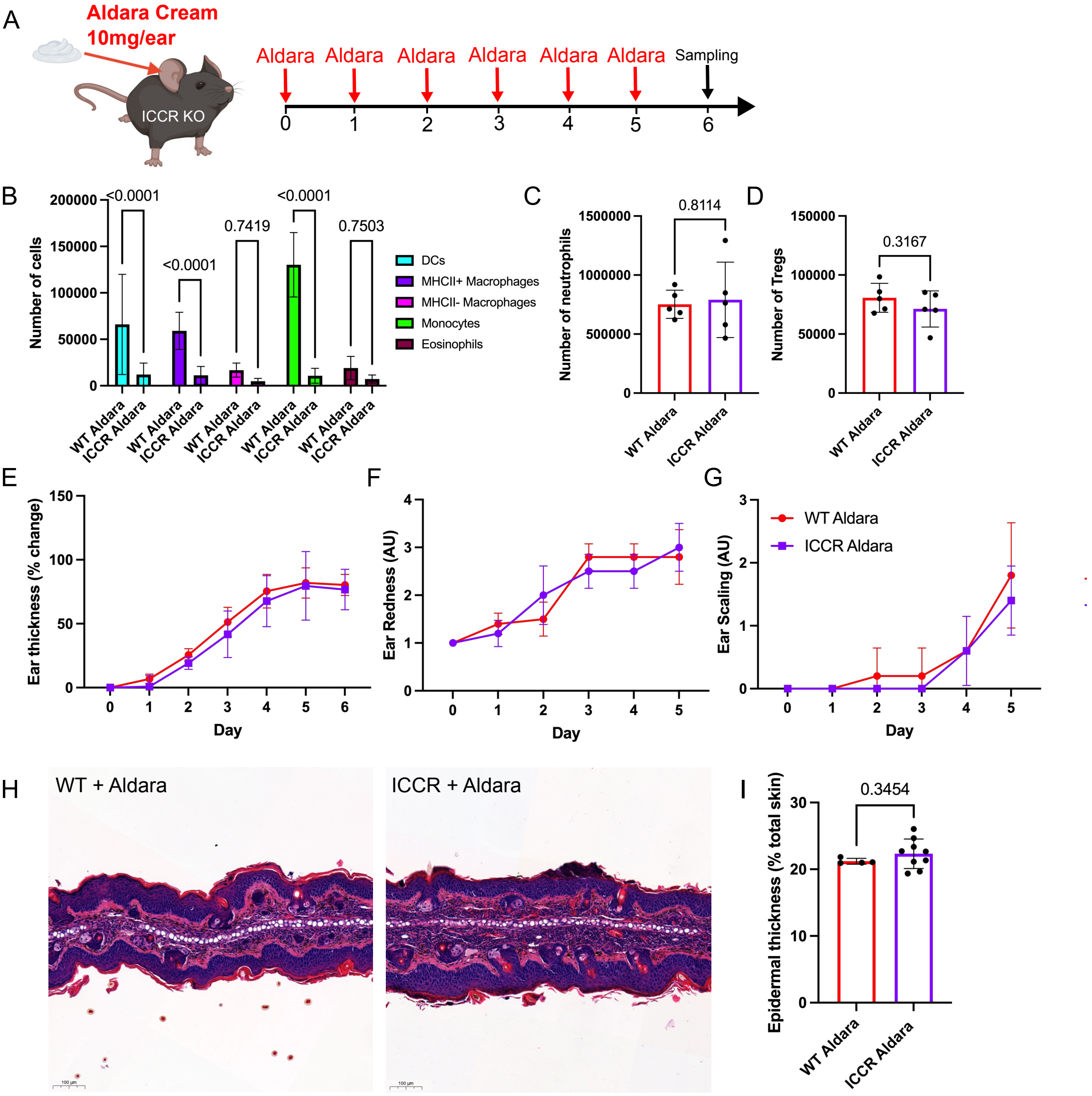


**Figure S6**. *Depletion of myeloid cells does not recapitulate the enhanced inflammation seen with heparanase inhibition during psoriasis-like skin inflammation.* ICCR KO mice lacking CCRs 1, 2, 3 and 5 as well as WT littermate controls were treated daily with topical application of Aldara cream to the ear pinnae for 6 days (**A**). Numbers of dendritic cells, macrophages, monocytes and eosinophils (**B**) as well as neutrophils (**C**) and Tregs (**D**) were quantified in skin by flow cytometry. Clinical readouts of inflammation were measured including ear thickness (**E**), redness (**F**) and scaling (**G**). Skin was sectioned and stained using haematoxylin and eosin (H&E) (**H**) and epidermal thickness measured and calculated as a % of total ear thickness (**I**). Histology scale bars in **H** represent 100μm. Data show 2 pooled experiments, except from **H** which shows representative images at 10x magnification. Data were analysed by one-way ANOVA and Tukey's multiple comparison test, except from **F and G** which are discontinuous variable and so were analysed by a non-parametric Kruskal-Wallis test with Dunn's multiple comparisons. **C, D and I** was analysed by unpaired t test. Data with no p values shown are not significant ($p>0.05$). Error bars represent mean ± SD.

#### Supplementary methods

**Table S1**. *Antibodies used for flow cytometry staining.*

| Antigen | Conjugate | Dilution | Clone | Supplier |
|---|---|---|---|---|
| CD11b | FITC | 1:300 | M1/70 | eBioscience |
| CD11c | BV605 | 1:800 | N418 | BioLegend |
| CD3 | APC-Cy7 | 1:100 | 17A2 | eBioscience |
| CD31 | BV711 | 1:200 | 390 | BD Biosciences |
| CD45 | BV510 | 1:300 | 30-F11 | BioLegend |
| F4/80 | BV786 | 1:500 | BM8 | BioLegend |
| Fc block (CD16/CD32) | Unconjugated | 1:1000 | 2.4G2 | BD Bioscences |
| Foxp3 * | BV421 | 1:100 | FJK-16s | eBioscience |
| **Heparan Sulfate #** | Biotin | 1:200 | F58-10E4 | Amsbio |
| Heparanase * | None (Rabbit IgG) | 1:200 | N/A | Proteintech |
| Interleukin-17* | PE-Cy7 | 1:100 | eBio1787 | eBioscience |
| Ly6C | PerCPCy5.5 | 1:500 | HK1.4 | BioLegend |
| Ly6G | AF700 | 1:200 | 1A8 | BioLegend |
| MHCII | BV650 | 1:500 | MS/114.15.2 | BioLegend |
| Siglec F | PEef610 | 1:400 | E50-2440 | BD Biosciences |
| Streptavidin | PEef610 | 1:200 | N/A | BioLegend |
| TCRβ | APC-Cy7 | 1:200 | H57-597 | Ebioscience |
| TCRγδ | APC | 1:300 | GL3 | eBioscience |
| Rabbit IgG | PE | 1:200 | Polyclonal | Rockland |

**Table S2**. *Antibodies for immunofluorescence staining.*

| Reagent | Conjugate | Dilution | Used to detect | Species | Clone | Supplier |
|---|---|---|---|---|---|---|
| Mouse anti-Heparan Sulfate | Unconjugated | 1:250 | Heparan Sulfate | Mouse | F58-10E4 | Amsbio |
| M.O.M biotinylated anti-Mouse IgG reagent | Biotin | 1:200 | Primary HS antibody | N/A | N/A | Vector |
| Streptavidin | NL-557 | 1:800 | Biotin on secondary HS antibody | N/A | N/A | R&D systems |
| Rabbit anti-CD31 | Unconjugated | 1:50 | Endothelial cells | Rabbit | DBV9E | Cell signaling Technology |
| Donkey anti-rabbit IgG | NL-637 | 1:100 | Primary CD31 antibody | Donkey | N/A | R&D systems |
